# Increased apoptotic priming of glioblastoma enables therapeutic targeting by BH3-mimetics

**DOI:** 10.1101/2021.06.13.448232

**Authors:** Anna L Koessinger, Dominik Koessinger, Kevin Kinch, Laura Martínez-Escardó, Nikki R Paul, Yassmin Elmasry, Gaurav Malviya, Catherine Cloix, Kirsteen J Campbell, Florian J Bock, Jim O’Prey, Katrina Stevenson, Colin Nixon, Mark R Jackson, Gabriel Ichim, William Stewart, Karen Blyth, Kevin M Ryan, Anthony J Chalmers, Jim C Norman, Stephen WG Tait

**Author notes:** To whom correspondence should be addressed: Stephen Tait CRUK Beatson Institute, University of Glasgow, Glasgow, G61 1BD, UK.

## Abstract

*IDH* wild-type glioblastoma (GBM) is the most prevalent malignant primary brain tumour in adults. GBM typically has a poor prognosis, mainly due to a lack of effective treatment options leading to tumour persistence or recurrence. Tackling this, we investigated the therapeutic potential of targeting anti-apoptotic BCL-2 proteins in GBM. Levels of anti- apoptotic BCL-xL and MCL-1 were consistently increased in GBM compared with non- malignant cells and tissue. Moreover, we found that relative to their differentiated counterparts, patient-derived GBM stem-like cells also displayed higher expression of anti- apoptotic BCL-2 family members. Surprisingly, high anti-apoptotic BCL-xL and MCL-1 expression correlated with heightened susceptibility of GBM to BCL-2 family protein- targeting BH3-mimetics. This is indicative of increased apoptotic priming. Indeed, GBM displayed an obligate requirement for MCL-1 expression in both tumour development and maintenance. Investigating this apoptotic sensitivity, we found that sequential inhibition of BCL-xL and MCL-1 led to robust anti-tumour responses *in vivo*, in the absence of overt toxicity. These data demonstrate that BCL-xL and MCL-1 pro-survival function is a fundamental prerequisite for GBM survival that can be therapeutically exploited by BH3- mimetics.

## Introduction

In adults, *IDH* wild-type glioblastoma *(IDHwt* GBM) is the most prevalent and malignant primary brain tumour (1, 2). Despite current multimodal treatment, comprising surgical resection with adjuvant radiotherapy and alkylating chemotherapy, the median survival in newly diagnosed patients remains poor at less than 12 months (3, 4). Resistance to conventional radio- and chemotherapy primarily emerges from persistent cancer stem cells, a tumourigenic subpopulation of GBM cells, consisting of heterogenous subclones and capable of self-renewal (5, 6). Therefore, targeting cells with stem-like capabilities is essential to develop effective treatment options and improve patient survival.

Treatment resistance can often be attributed to cells circumventing therapy-induced cell death. Apoptosis is an evolutionarily conserved type of cell death with broad ranging importance in biology (7). The intrinsic (mitochondrial) pathway of apoptosis is controlled by pro- and anti-apoptotic members of the B cell lymphoma 2 (BCL-2) family that regulate mitochondrial outer membrane integrity (8). During apoptosis, pro-apoptotic BCL-2 proteins cause mitochondrial outer membrane permeabilisation or MOMP. This leads to the release of mitochondrial intermembrane space proteins, including cytochrome *c*, that activate caspase proteases leading to apoptotic cell death (8).

Increased anti-apoptotic BCL-2 protein expression has been described in a wide range of solid cancers and is often linked with insensitivity to conventional chemotherapy (9–11). Recently, a new class of chemotherapeutics called BH3-mimetics have been developed that target pro-survival BCL-2 function, sensitising to cell death. BH3-mimetics have proven to be highly effective in haematologic malignancies. For instance, venetoclax (ABT-199), a BCL-2 targeted BH3-mimetic (12), is in clinical use for chronic lymphocytic leukaemia (CLL) (13) and acute myelogenous leukaemia (AML) (14, 15). CLL cells typically express high levels of anti-apoptotic BCL-2 protein. Nevertheless, the high intrinsic apoptotic sensitivity - also called apoptotic priming - of CLL renders it sensitive to venetoclax. For solid cancers, venetoclax is currently being tested in combination with conventional chemotherapeutic agents. The combination of venetoclax and tamoxifen has progressed to early phase clinical trials in patients with estrogen receptor positive (ER+), high BCL-2 expressing breast cancer (16). Other BH3-mimetics developed to target BCL-xL and MCL-1 have shown promising pre-clinical results in combination with inhibitors of MEK1/2 for solid cancers harbouring oncogenic mutations in the MAPK pathway (17–19). Approximately 10% of GBM carry a mutation of isocitrate dehydrogenase 1 (*IDH1*) (1), which has been linked with increased sensitivity to treatment with BH3-mimetics targeting BCL-xL (20). Furthermore, previous studies have proposed BCL-xL as a treatment target in combination with ionising radiation (21) and other chemotherapeutics (22) in GBM.

Because tumours retain characteristics of their tissue origins, brain derived glial cancers exhibit defined cellular hierarchies found in brain development and homeostasis (23–25). During central nervous system development, anti-apoptotic BCL-2 family proteins play a pivotal role in promoting cell survival (26, 27) while with adulthood the brain becomes refractive to apoptosis (28). Given this important role in cell survival, we hypothesised that GBM, while phenocopying the developing brain, might display similar anti-apoptotic survival dependencies. Indeed, we found increased levels of the major pro-survival proteins in GBM, specifically within the stem-cell enriched population. Surprisingly, high BCL-xL and MCL-1 expression correlates with increased apoptotic sensitivity, demonstrating that GBM stem-like cells are primed for apoptosis. Exploiting this, we found that sequential dosing of BCL-xL and MCL-1 targeting BH3-mimetics enables effective treatment responses both, *in vitro* and *in vivo*. This could offer a therapeutically tractable approach for patients with *IDHwt* GBM.

## Results

### High anti-apoptotic BCL-2 family protein expression correlates with increased BH3-mimetic sensitivity in GBM

Cancer stem cells are proposed to give rise to GBM and contribute to therapeutic resistance (23). We therefore sought to assess the apoptotic sensitivity of GBM stem-like cells (GSC) by treating them with BH3-mimetics with selectivity for BCL-2, BCL-xL and MCL-1. For this purpose, we used a panel of patient-derived *IDHwt* GSC, cultured under conditions to maintain their tumour specific phenotype and stem cell properties (29, 30). Cell viability was measured using IncuCyte live-cell imaging and SYTOX Green exclusion. Importantly, three cell lines (G1, G7 and R24 GSC) were sensitive to A-1331852, a selective BCL-xL antagonist (31) whereas two cell lines (R9 and R15 GSC) displayed sensitivity to S63845, a potent and selective MCL-1 inhibitor (32) (**Figure 1A,B**). Moreover, the commonly used GBM cell line U87MG displayed increased sensitivity to BCL-xL inhibition when cultured under stem cell-enriching conditions. One cell line (E2 GSC) was resistant to all single agent treatments. Treatment with venetoclax (ABT-199), a BCL-2 specific inhibitor, induced no more than 26% cell death in any GSC and therefore was comparably inefficient. Collectively, these data show that the majority of tested GSC display survival dependence on anti- apoptotic BCL-2 family function.

**Figure 1.**
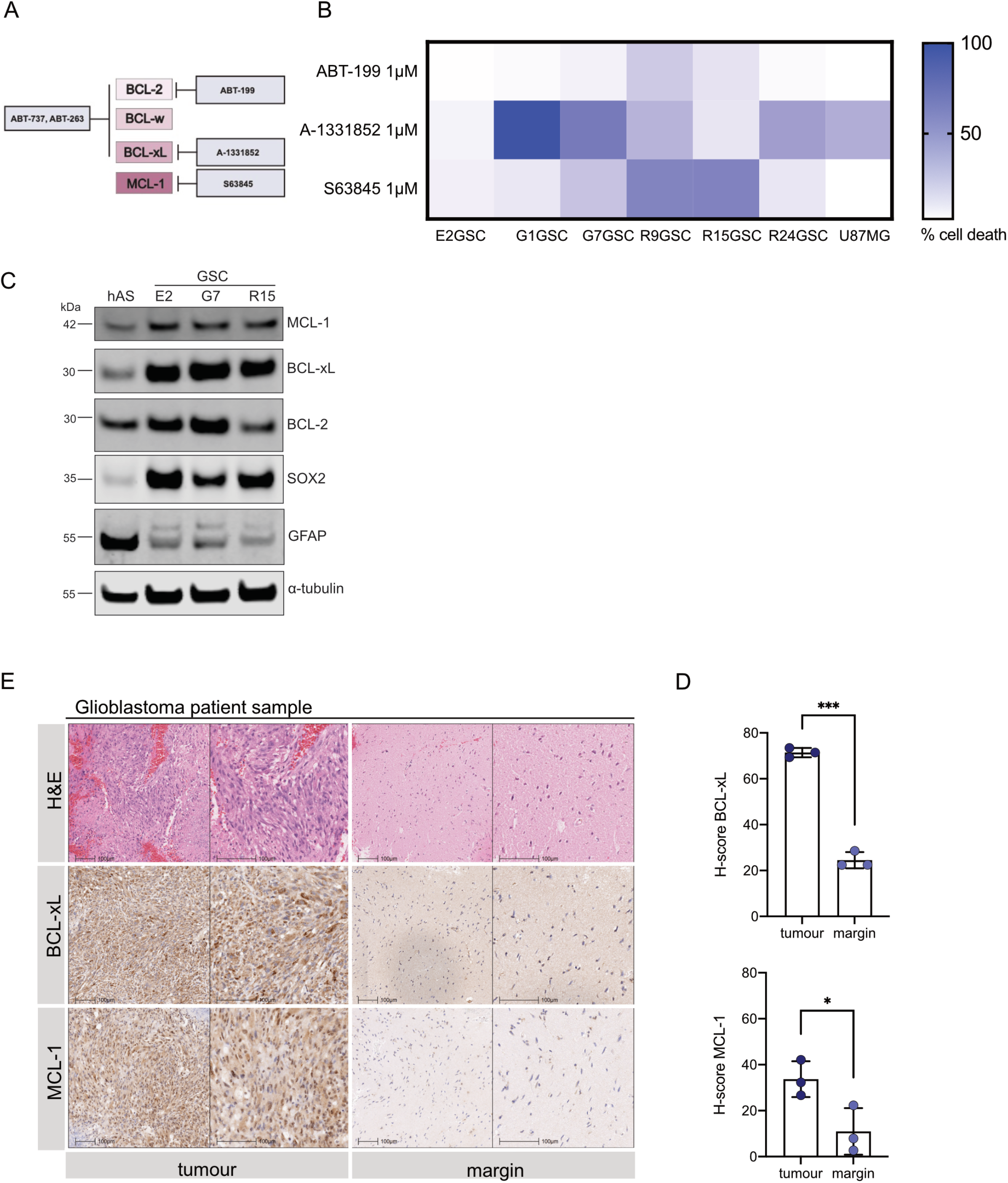
High anti-apoptotic BCL-2 family protein expression correlates with increased BH3-mimetic sensitivity in GBM. (**A**) Schematic overview of BH3-mimetic drugs used and their respective targets (**B**) Panel of six GSC cell lines and the human primary GBM cell line U87MG were treated with indicated drugs for 24 to 48 hours and analysed for cell viability using an IncuCyte imager and SYTOX Green exclusion. Results are presented as heatmap. Percentage cell death was calculated by normalising against maximal cell death (treatment with 1µM Actinomycin D, 10µM ABT- 737 and 1µM S6384), n=3 independent experiments per cell line. (**C**) Immunoblot of BCL-2 family proteins, cell-line specific neural stem cell marker SOX2 and astrocyte lineage differentiation marker GFAP in human astrocytes and patient-derived GSC. α-tubulin served as loading control. Representative image from n=3 independent experiments. (**D,E**) Matched tumour and margin specimens were obtained from three patients diagnosed with GBM and stained for haematoxylin and eosin (H&E), BCL-xL and MCL-1 IHC (representative images of one case shown). H-score (analysis of intracellular BCL-xL and MCL-1 expression) was determined using automated analysis with Halo. Error bars represent mean +/-SD (***p=0.0002, *p=0.0404) Welch’s test. Further representative images in lower magnifications are shown in Supplementary Figure 1C.

We next used immunoblotting to determine if the individual apoptotic sensitivity of the patient-derived GSC corresponded to anti-apoptotic BCL-2 protein expression. In comparison to human differentiated astrocytes, all GSC exhibited higher expression of BCL- xL and MCL-1 and partially higher expression of BCL-2 (**Figure 1C**). Consistent with their origins, GSC expressed higher levels of neural stem cell marker SOX2 (33), while cell lineage specific GFAP was more abundant in astrocytes (34). Subsequently, we investigated whether anti-apoptotic BCL-2 protein expression also differed in GBM tumours and adjacent brain tissue. Using immunohistochemistry (IHC) we compared BCL-xL and MCL-1 expression in matched specimens of three patients diagnosed with GBM. In all cases, MCL- 1 and BCL-xL were increased in the tumour cores compared to the predominantly non- tumorous margins (**Figure 1D,E**). Extending this analysis, we determined MCL-1 and BCL-xL mRNA expression in different glioma subtypes and normal brain tissue using the publicly available REMBRANDT database. In line with our IHC analysis, *BCL-xL* and *MCL-1* mRNA levels were more expressed in GBM compared with lower grade gliomas and normal brain tissue (**Supplementary Figure 1A,B**). These data demonstrate specific sensitivities of patient-derived GSC to individual BH3-mimetics and increased expression of anti-apoptotic BCL-2 proteins in both primary GBM tumour tissues and GSCs.

### Anti-apoptotic MCL-1 is required for the growth and survival of GBM

While it has previously been shown that GBM tumoursphere formation is promoted by high BCL-xL expression (35), little is known about the role of MCL-1 on GBM growth and maintenance. To explore the importance of MCL-1 in GBM formation and growth *in vivo*, we selected a tumourigenic cell line (G7 GSC) and deleted *MCL-1* using CRISPR/Cas9 genome editing. Western blot analysis confirmed efficient *MCL-1* deletion, while the expression of BCL-xL, SOX2 and GFAP was increased in the MCL-1^CRISPR^ cells (**Supplementary Figure 2A**). MCL-1^CRISPR^ GSC were found to proliferate at the same rate and retained a similar capability to form neurospheres as their vector^CRISPR^ counterparts (**Figure 2A, Supplementary Figure 2B**). We next investigated whether MCL-1 was required for tumourigenesis *in vivo*. iRFP-labelled vector^CRISPR^ and MCL-1^CRISPR^ G7 GSC were orthotopically injected in CD-1 nude mice and tumour growth was monitored with cranial magnetic resonance imaging (MRI) and iRFP signal detection (36). We observed a substantial impairment of tumour growth in MCL-1 deleted tumours (**Figure 2B,C; Supplementary Figure 2C**) that was reflected in the significantly prolonged survival of these mice (**Figure 2D**). Importantly, IHC analysis of the end-stage tumours revealed an outgrowth of MCL-1 proficient tumour cells in the MCL-1^CRISPR^ xenografts (**Supplementary Figure 2D**). In contrast to the *in vitro* analysis, these data reveal a key role for MCL-1 in initiation and growth of GBM *in vivo* and identify MCL-1 as a promising therapeutic target. Our results also support an important pro-survival role for anti-apoptotic MCL-1 in GBM.

**Figure 2.**
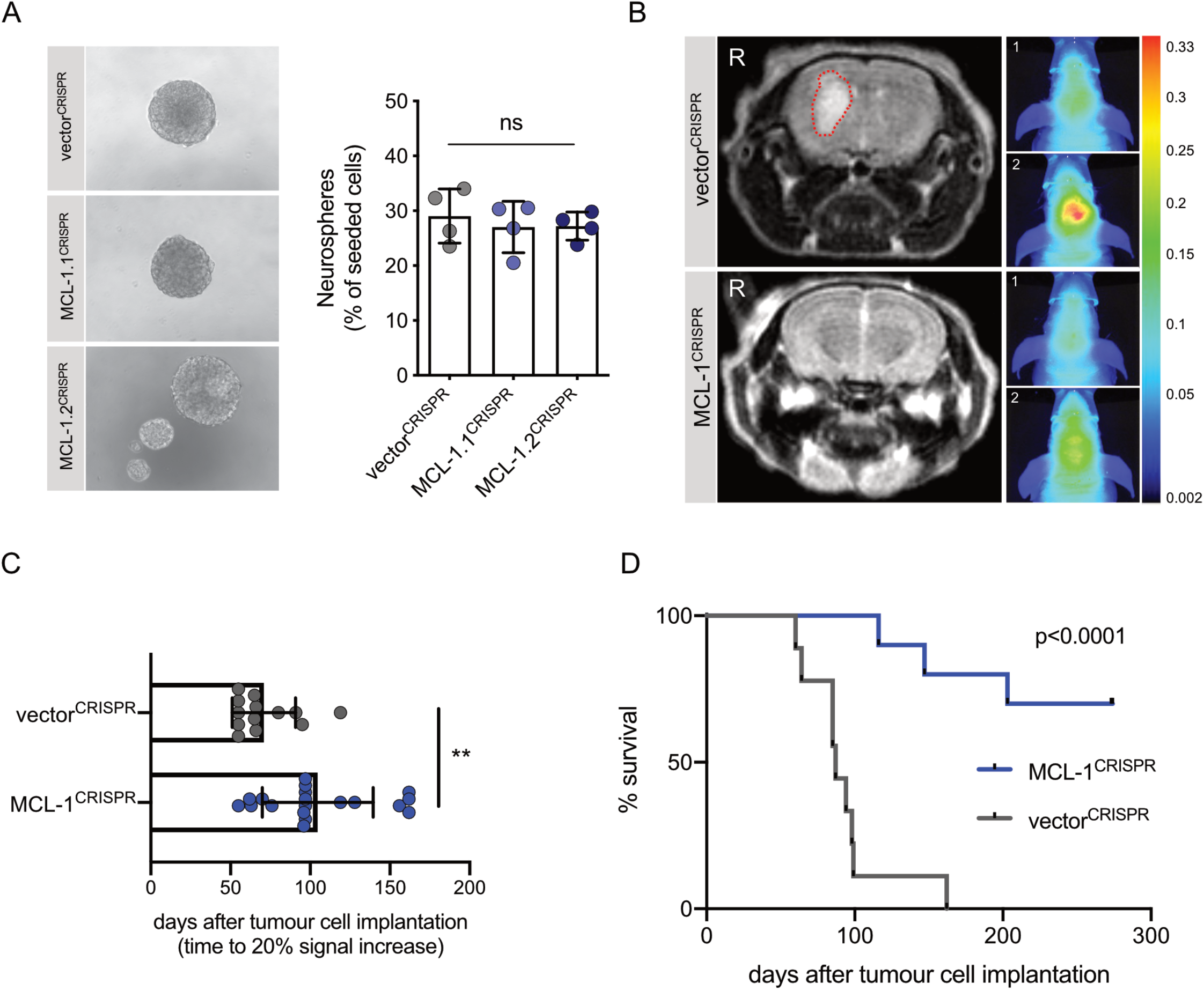
Anti-apoptotic MCL-1 is required for the growth and survival of GBM. (**A**) Representative images of neurosphere growth from G7 GSC vector^CRISPR^ (upper panel), MCL-1.1^CRISPR^ (middle panel) MCL-1.2^CRISPR^ (lower panel), respectively. Quantification of neurosphere formation capacity by G7 GSC vector^CRISPR^ vs. MCL-1.1^CRISPR^ or MCL-1.2^CRISPR^. Error bars represent mean +/-SD from n=4 independent experiments (p=0.8146, ns, nonsignificant) Welch’s test. (**B**) Representative images of brain MRI scans (tumour indicated by red dashed line) next to corresponding pseudocolour representations of iRFP signal of mice bearing iRFP tagged G7 GSC vector^CRISPR^ (upper panel) and MCL-1.2^CRISPR^ (lower panel) xenografts, respectively. iRFP signal was detected by PEARL scans (700nm channel) (1) at week 8 and (2) at week 20 (vector^CRISPR^) or week 36 (MCL1^CRISPR^) post injection. (**C**) Quantification of time to 20% iRFP signal increase of G7 vector^CRISPR^ n=13 vs. G7 MCL-1^CRISPR^ tumours n=19, compared to 4 weeks post injection (baseline signal). Error bars represent mean +/-SD (**p=0.0013) Mann-Whitney test. (**D**) Kaplan-Meier survival graph of mice with orthotopic xenografts of G7 GSC iRFP vector^CRISPR^ n=9 (median survival 87 days) vs. MCL-1^CRISPR^ tumours n=10 (median survival undefined) post tumour cell implantation (p<0.0001) Log-rank (Mantel-Cox) test.

### GSC display increased apoptotic priming and can be effectively killed by dual BCL-xL, MCL- 1 inhibition

Currently, no treatment regimen is able to achieve long-time remission of GBM, with tumours inevitably developing resistance to treatment and recurring, eventually leading to patient death (37). Anti-apoptotic BCL-2 family members have overlapping binding affinities for several pro-apoptotic BH3-only proteins (8). We asked whether GBM might circumvent single inhibitor treatment by compensatory upregulation of untargeted anti-apoptotic proteins. Indeed, upon treatment with BCL-xL inhibitor A-1331852 or navitoclax (ABT-737), an inhibitor of BCL-xL, BCL-2 and BCL-w, we found that levels of MCL-1 protein were increased in surviving GSC (**Figure 3A, Supplementary Figure 3A**). We reasoned that GBM might counteract drug-induced neutralisation of BCL-xL function via increased MCL-1 expression. To address this, we treated MCL-1^CRISPR^ G7 and R24 GSC with the BCL-xL inhibitors A-1331852 or ABT-263. Cell viability was measured by live-cell IncuCyte imaging with Sytox Green exclusion or in a clonogenic survival assay. In all cases, MCL-1 deletion significantly increased cellular sensitivity to the BCL-xL specific antagonist and navitoclax (**Figure 3B, Supplementary Figure 3B**). Similarly, dual inhibition of BCL-xL and MCL-1 with A-1331852 and S63845 displayed a substantial combinatorial effect resulting in up to 100% cell death across a range of GSC, determined in both short term cell viability assays and long-term clonogenic survival assays (**Figure 3C,D; Supplementary Figure 3C,D**). This effect was observed even at 10-fold decreased doses compared to effective single treatment. Verifying on-target engagement of mitochondrial apoptosis, combined MCL-1 and BCL-xL inhibition led to Caspase 3 and PARP-1 cleavage as well as cell death in a BAK, BAX and caspase-dependent manner (**Figure 3E, Supplementary Figures 3E-F**).

**Figure 3.**
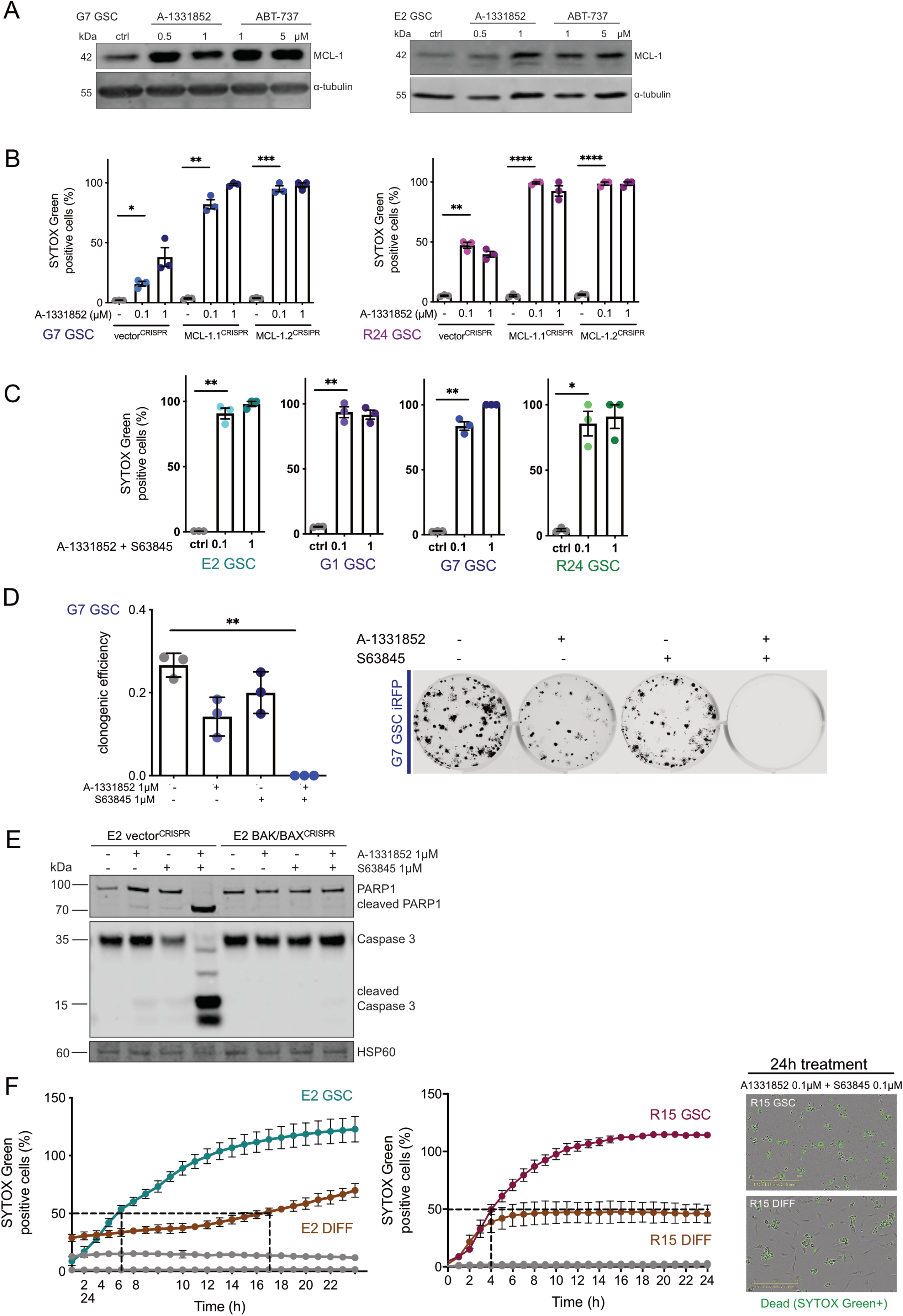
GSC display increased apoptotic priming and can be effectively killed by dual BCL-xL, MCL-1 inhibition. (**A**) G7 and E2 GSC were treated with DMSO (ctrl), A-1331852 or ABT-737 as indicated for 16 or 24 hours, respectively, harvested and protein expression was analysed by immunoblot. α-tubulin served as loading control. Representative image from n=3 independent experiments shown. (**B**) G7 or R24 GSC vector^CRISPR^ vs. MCL1.1^CRISPR^ and MCL1.2^CRISPR^ were treated with A-1331852 for 24 hours and analysed for cell viability using an IncuCyte imager and SYTOX Green exclusion. Percentage cell death was calculated by normalising against maximal cell death verified by visual inspection. Error bars represent mean +/-SEM from n=3 independent experiments. (G7: *p=0.0218, **p=0.002, ***p=0.0006) (R24: **p=0.0028, ****p<0.0001) Welch’s test. (**C**) E2, G1, G7, R24 GSC were treated with a combination of A-1331852 and S63845 in indicated concentrations for 24 hours and analysed for cell viability using an IncuCyte imager and SYTOX Green exclusion. Percentage cell death was calculated by normalising against maximal cell death as described in Figure 1B. Error bars represent mean +/-SEM from n=3 independent experiments. (E2 **p=0.0021, G1 **p=0.0022, G7 **p=0.0016, R24 *p=0.0123) Welch’s test. (**D**) Clonogenic survival assay of G7 GSC iRFP treated with indicated drugs 16 hours after plating 250 cells per well. Colonies counted manually after 14 days. Error bars represent mean +/-SD from n=3 independent experiments (**p=0.0038) Welch’s test. Representative images of a replicate in one independent repeat scanned on LICOR imager. (**E**) E2 GSC vector^CRISPR^ and BAK/BAX^CRISPR^ were treated as indicated for 2 hours, harvested and protein expression was analysed by immunoblot. HSP60 served as loading control. Representative image from n=3 independent experiments. (**F**) E2 and R15 GSC and paired DIFF cells were treated either with DMSO (grey) or a combination of A-1331852 and S63845 (both 0.1 μM) for 24 hours and analysed for cell viability using an IncuCyte imager and SYTOX Green exclusion. Error bars represent mean +/-SEM from n=3 independent experiments. Representative IncuCyte images 24 hours after treatment are shown.

Sensitivity of GBM cells to chemotherapy and ionising radiation inversely correlates with tumour cell stemness (6, 38, 39). We therefore hypothesised that the differentiated counterparts (DIFF) of the patient-derived GBM stem-like cells may be more sensitive towards BH3-mimetic treatment. To ensure comparable culture conditions for GSC and DIFF, we conducted these experiments using 1% FCS containing Ad-DMEM medium during the experimental procedure. Cells were treated with A-1331852 and S63845 to inhibit BCL-xL and MCL-1 respectively and cell viability measured by live-cell IncuCyte imaging and Sytox Green exclusion. Following treatment with single BH3-mimetic, we found that cell viability was largely comparable for DIFF and GSC (**Supplementary Figure 3G**).

Surprisingly, following dual MCL-1 and BCL-xL inhibition, E2 and R15 GSC were more sensitive than DIFF cells; while >50% cell death was observed within about 5 hours in GSC, it was not observed in DIFF cells until 16 hours (**Figure 3F**). Moreover, 100% cell death was not achieved in either of the DIFF cell lines. Together, these data suggest that GSCs are more primed for apoptotic cell death than DIFF cells. To investigate this further, we compared expression of pro- and anti-apoptotic BCL-2 proteins in the paired cell lines.

Although more sensitive to apoptosis, GSCs displayed higher levels of anti-apoptotic BCL-2 proteins, BCL2, BCL-xL and MCL-1 than their differentiated counterparts (**Supplementary Figure 3H**). In summary, these data indicate that GSC can display increased apoptotic priming and reveal potent cytotoxic effects of dual-targeting BCL-xL and MCL-1.

### TrkB signalling regulates sensitivity of GSC to anti-apoptotic treatment

We next sought to explore the differential priming between GSC and their isogenic differentiated counterparts. To this end, bulk RNA sequencing data from E2, G7 GSC and their DIFF counterparts was analysed (30). Consistent with enrichment of GSC, RNAseq analysis revealed increased levels of CD34, a surface glycoprotein, first described as marker for haematopoietic progenitor cells (40). Interestingly, high expression of *NTRK2* mRNA was detected in both GSC (**Figure 4A**). This finding was validated in E2 and G7 GSC as well as R15 and R24 GSC via immunoblotting (**Figure 4B, Supplementary Figure 4A**). *NTRK2* the gene coding for the tropomycin receptor kinase B (TrkB) is primarily known for its function in neurodevelopment inducing downstream signalling upon binding of brain-derived neurotrophic factor (BDNF) (41). Recently, Wang and colleagues have reported a role for TrkB-expressing cancer stem cells in GBM progression in response to BDNF stimulation by differentiated tumour cells (42). As TrkB-mediated activation of MAPK and PI3K-AKT signalling is generally associated with cell survival (43), we hypothesised that BDNF-mediated TrkB stimulation might enable GSC to evade cell death. Unexpectedly, following stimulation of GSC with BDNF or 7,8-dihydroxyflavone (7,8-DHF), a specific TrkB agonist, we found that GSC were further sensitised to cell death following treatment with BH3- mimetics targeting BCL-xL and or MCL-1 (**Figure 4C, Supplementary Figure 4B**). This sensitising effect was not observed in the DIFF cells (**Figure 4D**). DIFF cells express significantly lower levels of the TrkB receptor and therefore prove to be comparably unresponsive to BDNF stimulation (**Supplementary Figure 4C**). BDNF-induced TrkB phosphorylation also led to increased BCL-xL protein expression, alongside stabilisation of the BIM protein downstream of MAPK signalling, independently of BCL-xL and MCL-1 inhibition (**Figure 4E**). These data demonstrate a key role for BDNF-TrkB signalling in the increased apoptotic priming of GSCs.

**Figure 4.**
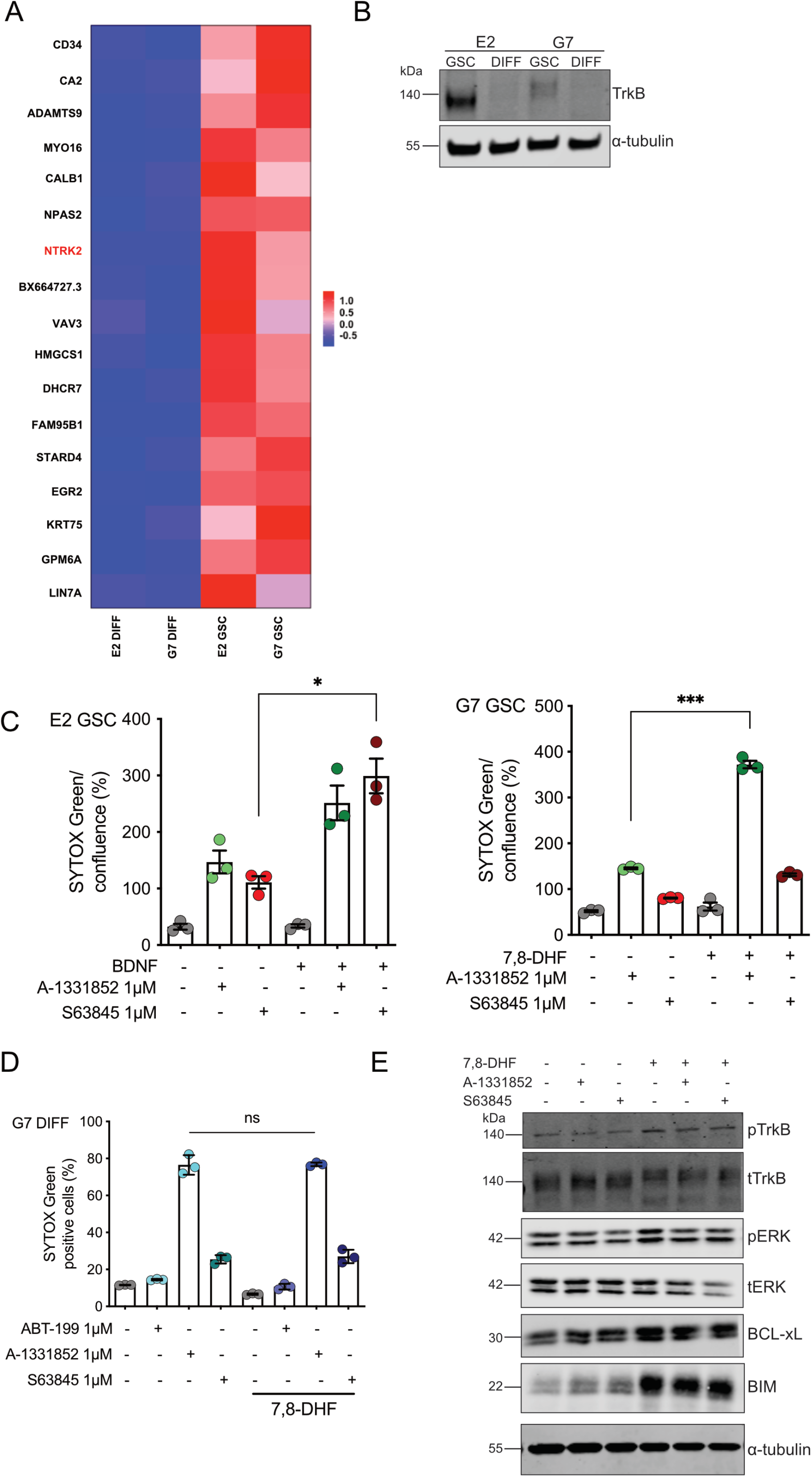
TrkB signalling regulates sensitivity of GSC to anti-apoptotic treatment. (**A**) Most differentially expressed genes in RNAseq analysis of E2 and G7 GSC vs. E2 and G7 DIFF. *NTRK2* codes for TrkB. (**B**) Immunoblot of TrkB in E2 and G7 GSC compared with paired DIFF cells. α-tubulin served as loading control. Representative image from n=2 independent experiments. (**C**) E2 and G7 and GSC treated with BDNF (100ng/mL) or 7,8- DHF (20μg/mL) +/- A-1331852 1μM and S63845 1μM for 24 hours (after a 24-hour starvation period in 1% glutamine containing DMEM/F12) and analysed for cell viability using an IncuCyte imager and SYTOX Green exclusion. Error bars represent mean +/-SEM from one of n=3 independent experiments (E2 *p=0.0166, G7 ***p=0.008) Welch’s test. (**D**) G7 DIFF treated as described in Figure 4C and analysed for cell viability using an IncuCyte imager and SYTOX Green exclusion. Error bars represent mean +/-SEM from n=3 independent experiments (ns, p=0.9284). (**E**) G7 GSC were treated with 7,8-DHF (20μg/mL) +/- A-1331852 1μM and S63845 1μM for 1 hour, harvested and protein expression was analysed by immunoblot. α-tubulin served as loading control. Representative image from n=3 independent experiments.

### Combined BCL-xL and MCL-1 inhibition causes apoptosis in human GBM ex vivo

Current *in vitro* methodologies fail to recapitulate important aspects of the brain microenvironment and tissue context. Given this, we sought to use a more physiologically relevant model to investigate functional responses to BCL-xL and MCL-1 inhibition in GBM. For this purpose, we developed an assay tailored to the use of freshly resected human GBM to be cultured *ex vivo* as tissue slices that could be readily exposed to candidate drugs (experimental setup illustrated in **Supplementary Figure 5A**). All three patients included in the study were diagnosed with *IDHwt* GBM. Tissue slices were treated for 72 hours in total. In all cases, we found that a combined therapy with A-1331852 and S63845 (BCL-xL and MCL-1 inhibition) significantly reduced tumour cell count compared with single drug treatment or control (**Figure 5A-D, Supplementary Figure 5B-D**). Moreover, dual treatment induced a significant reduction of cell proliferation (Ki67 IHC) and amount of SOX2 positive tumour cells (**Figure 5E,F, Supplementary Figure 5E,F**), while Caspase 3 cleavage was increased. This data indicates that the dual treatment efficiently targets GBM stem-like cells *ex vivo*. Importantly, the integrity of the brain tissue and vasculature was maintained (**Supplementary Figure 5G**).

**Figure 5.**
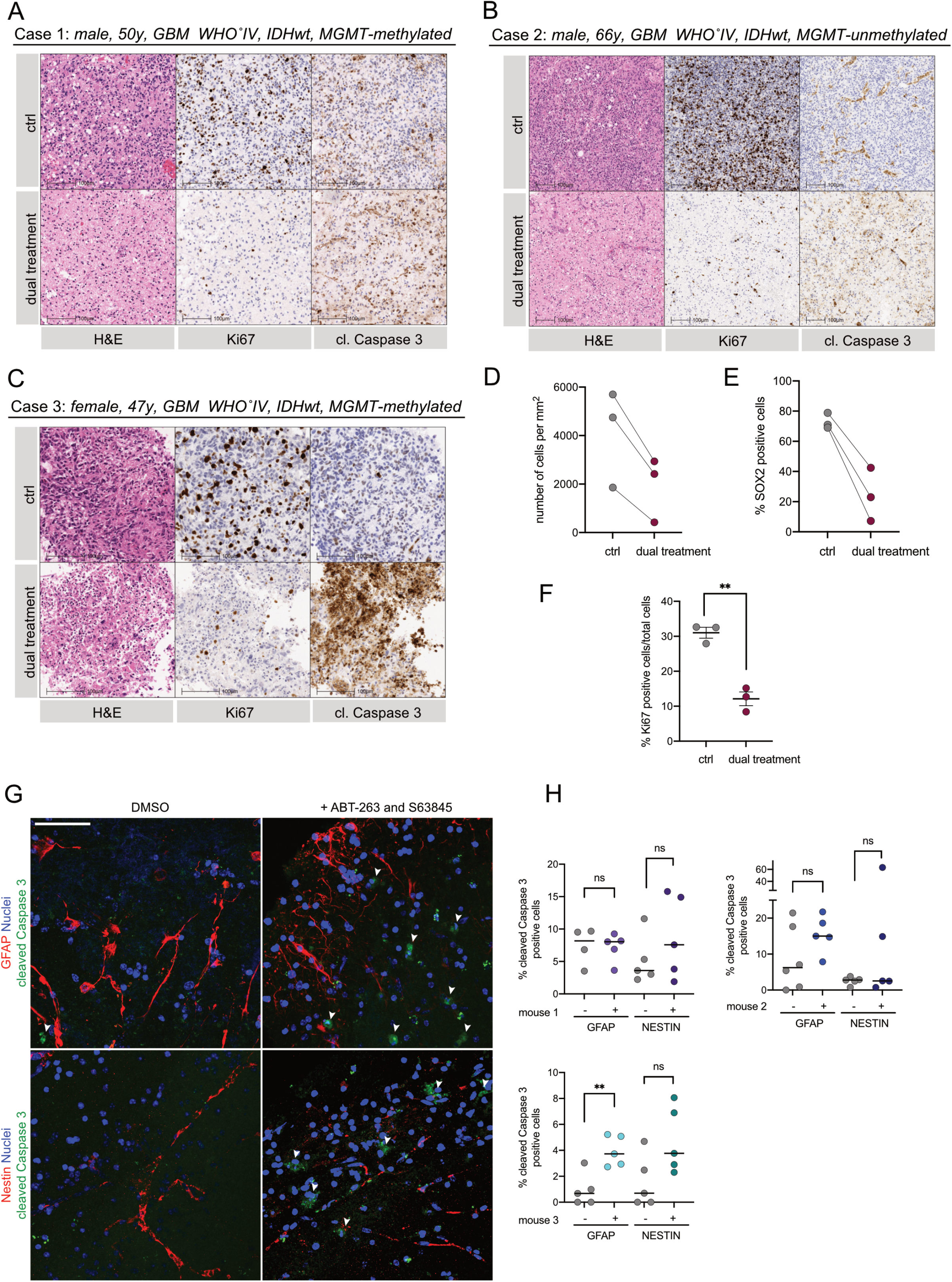
Combined BCL-xL and MCL-1 inhibition causes apoptosis in human GBM *ex vivo*. (**A**-**C**) Representative images of H&E, Ki67 and cleaved Caspase 3 IHC of three patients diagnosed with *IDHwt* GBM (case 1-3; ctrl=DMSO, dual treatment=A-1331852 1μM + S63845 1μM) (**D-F**) Quantification of cellularity, percentage Ki67 and SOX2 positive cells/total cell count in all three cases treated with the drugs described in (A-C) for 72 hours. Error bars represent mean +/-SEM (**p=0.0021) Welch’s test. (**G**) Representative images of IF staining of GFAP, NESTIN (both red) and cleaved Caspase 3 (green) in the subventricular zone (SVZ) of mouse brain slices, cultured and treated with DMSO or a combination of ABT-263 5μM and S63845 2μM for 24 hours, counterstaining with DAPI (blue). (**H**) Quantification of cleaved Caspase 3 positive cells normalised to nuclear count in single mice. Single dots represent analysed images. Error bars represent median (mouse 1: p=0.9198, p=0.3086; mouse 2: p=0.149, p=0.2974; mouse 3: **p=0.0044, p=0.0593; ns, nonsignificant) Welch’s test. Scale bars = 50µm.

In recent years, selective MCL-1 and BCL-xL inhibitors have been developed that show effective *in vivo* potency (31, 32). However, systemic exposure to both inhibitors is limited due to its combined toxicity (44). Because the blood-brain-barrier is only permissible to certain drugs, local or intrathecal drug application (45) might allow to circumvent systemic side effects in a clinical setting. To explore potential toxicities to resident brain cells, we obtained brain slices from 11-week old adult mice and exposed them to dual treatment with the indicated BH3-mimetics. In regions of the subventricular zone, an important neural stem cell niche, we detected only a moderate increase of cleaved Caspase 3 in a fraction of glial and neural progenitor cells (GFAP and NESTIN IF stain) (**Figure 5G,H**). In summary, these data suggest that dual targeting of BCL-xL and MCL-1 may provide a novel therapeutic approach to treat GBM.

### Priming with BCL-xL inhibition renders GSC vulnerable to MCL-1 inhibition, promoting tumour regression in vivo

Given the potency of joint BCL-xL/MCL-1 inhibition in our *in vitro* and *ex vivo* findings, we sought to maximise this combinatorial effect whilst mitigating possible systemic toxicity. To address this, we investigated which pro-apoptotic proteins are involved in regulating intrinsic apoptosis in GBM. Upon single agent treatment with A-1331852, we observed upregulation of the BH3-only protein BIM as well as anti-apoptotic MCL-1. This was seen in both control and BAX/BAK deficient GSC (**Figure 6A**). BIM is an important BH3-only protein in the canonical apoptotic pathway where it functions by regulating both BCL-xL and MCL-1 mediated cell death responses (46). We hypothesised that MCL-1 might bind and neutralise BIM that is released by the A-1331852 complexing to BCL-xL. Accordingly, immunoprecipitation of MCL-1 following treatment with A-1331852 revealed increased binding of BIM to MCL-1 (**Figure 6B**). To explore whether this mechanism could be therapeutically exploited, we questioned whether BCL-xL inhibition would render GSC more sensitive to subsequent MCL-1 inhibition. GSC were treated with a BH3-mimetic targeting either BCL-xL or MCL-1 for up to 48 hours followed by a washout and 24 hours treatment pause. Subsequently the complementary inhibitor was applied for up to 48 hours. Whereas prior inhibition with MCL-1 inhibitor failed to sensitise the cells to BCL-xL inhibition, pre-treatment with the BCL-xL inhibitor substantially increased the susceptibility of GSC to subsequent MCL-1 inhibition (**Figure 6 C,D**). To further investigate the relevance of BIM in mediating apoptosis following BCL-xL and MCL-1 inhibition, we deleted BIM by CRISPR/Cas9 genome editing (**Supplementary Figure 6A**). Using IncuCyte live cell imaging and Sytox Green exclusion to detect cell death under treatment we observed that knockout of BIM did not impede the sensitivity of G7 GSC to concurrent dual BCL-xL and MCL-1 inhibition (**Supplementary Figure 6B**). However, after pre-treatment with BCL-xL inhibitor A-1331852 G7 BIM^CRISPR^ GSC were less primed for following MCL-1 inhibition compared to their vector^CRISPR^ counterparts (**Figure 6E**). These results indicate that Bcl-xL- inhibition mediates sensitisation of GSC to subsequent MCL-1 neutralisation via pro-apoptotic BIM.

**Figure 6.**
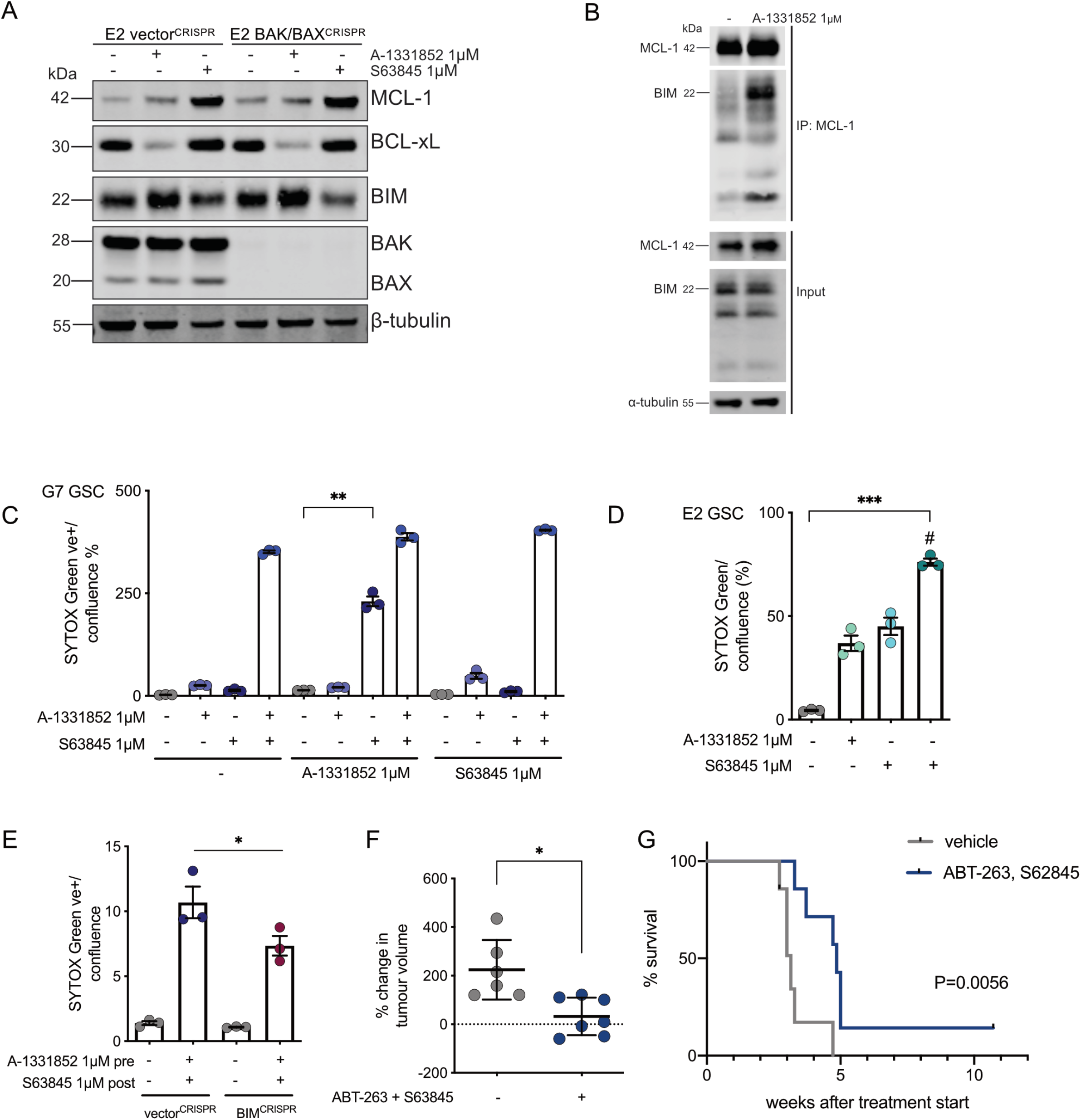
Priming with BCL-xL inhibition renders GSC vulnerable to MCL-1 inhibition, promoting tumour regression *in vivo*. (**A**) E2 GSC vector^CRISPR^ and BAK/BAX^CRISPR^ were treated with DMSO (-), A-1331852 and S63845 as indicated for 24 hours, harvested and protein expression was analysed by immunoblot. Representative image from n=2 independent experiments. β-tubulin served as loading control. (**B**) Direct binding interactions between MCL-1 and BIM were immunoprecipitated and interacting proteins were detected by western blot in G7 GSC treated for 16 hours with A-1331852 (Input=total cell lysate, IP=immunoprecipitated fraction). Representative image from n=2 independent experiments. (**C**) G7 GSC were treated with DMSO, A-1331852 or S63852 1μM for 48 hours, followed by 24 hours drug washout with exchange to fresh medium and treatment with indicated drugs for 24 hours. For cell viability analysis, IncuCyte imager and SYTOX Green exclusion was used. Error bars represent mean +/-SEM from n=3 independent experiments, (**p=0.003) Welch’s test. (**D**) E2 GSC were pre-treated with DMSO or A-1331852 1µM (#) for 24 hours, followed by 24 hours drug washout with exchange to fresh medium and treatment with indicated drugs for 48 hours. For cell viability analysis, IncuCyte imager and SYTOX Green exclusion was used. Error bars represent mean +/-SEM from n=3 independent experiments, (***p=0.0004) Welch’s test. (**E**) G7 GSC vector^CRISPR^ and BIM^CRISPR^ were treated for 48 hours with A-1331852 1µM, followed by 24 hours drug washout with exchange to fresh medium and treatment with S63845 1µM for 24 hours. For cell viability analysis, IncuCyte imager and SYTOX Green exclusion was used. Error bars represent mean +/-SEM from n=3 independent experiments, (*p=0.0132) Welch’s test. (**F**) Percent U87MG subcutaneous tumour volume change at the end of 2 weeks alternating treatment with ABT-263, followed by S63845, relative to tumour size at start. Treatment commenced when tumours were >5mm diameter. n=6 vehicle treated (grey dots) and n=7 drug treated (blue dots). Bars are mean +/- SD (*p=0.0104) Welch’s test. (**G**) Kaplan-Meier survival analysis of U87 vehicle treated (grey line, n=6, median survival 22 days) vs. U87 drug treated (blue line, n=7, median survival 34 days) since treatment start (**p=0.0056) Log-rank (Mantel-Cox) test.

Finally, we aimed to investigate the potency of alternating BH3-mimetic treatments in GBM *in vivo*. Due to the poor blood-brain-barrier penetrance of ABT-263, we chose a subcutaneous model. As the GSC used in our study do not grow as subcutaneous xenografts we explored whether human U87MG respond to combined and alternating inhibition of BCL-xL and MCL-1 in like manner to GSC. Indeed, we found that dual inhibition of BCL-xL and MCL-1 induced substantial cell death in U87MG neurospheres (**Supplementary Figure 6C**). Testing different treatment regimens using a clonogenic survival assay, we found a profound decrease in colony formation upon alternating treatment with ABT-263 and S63845 (**Supplementary Figure 6D**). In our *in vivo* cohort, mice were treated with vehicle or alternating therapy with ABT-263 and S63845 every 48 hours for two weeks upon tumour establishment (treatment schematic illustrated in **Supplementary Figure 6E**). Compared to mice receiving vehicle control, mice treated with the sequential therapy showed significant attenuation and/or regression of tumours (**Figure 6F**). Most importantly, we observed significantly improved survival in mice treated with ABT-263, followed by S63845 (**Figure 6G**). One mouse had a complete tumour regression after sequential treatment with no reoccurrence of the subcutaneous tumour over an 8-week follow up period. With this treatment schedule no significant weight loss (**Supplementary Figure 6F**) or signs of neurological deficits were detected in mice. Histopathological analysis of tumours reaching clinical end point showed a higher prevalence of a large central necrotic areas in within the treatment cohort compared to vehicle control (83% vs. 33%) (**Supplementary Figure 6G**). Collectively, these data demonstrate the therapeutic potential of sequential BCL-xL and MCL-1 inhibition in GBM.

## Discussion

Largely due to a dearth of effective treatment options, GBM patients have a dismal prognosis (37). Addressing this, we investigated the therapeutic potential of targeting anti-apoptotic BCL-2 proteins in GBM. Our analysis revealed high expression of anti-apoptotic BCL-xL and MCL-1 in GBM. Moreover, we also observed increased expression of BCL-xL and MCL-1 in GBM stem-like cells - a population of cells that are key for GBM development and treatment resistance *in vivo* (5, 6). Rather than promoting apoptotic resistance, elevated anti-apoptotic BCL-xL and MCL-1 expression in GSC compared to isogenic DIFF correlated with increased susceptibility to targeted inhibition using BH3-mimetics. This indicates that GSC are inherently primed for apoptosis. Exploiting this, we found that GBM were sensitive to BH3-mimetics targeting either MCL-1 or BCL-xL. Crucially, alternating dosing with BCL-xL followed by MCL-1 specific BH3-mimetics, led to durable treatment responses with preceding BCL-xL inhibition sensitising to MCL-1 inhibition *in vivo*. These data highlight the therapeutic potential of targeting BCL-xL and MCL-1 in GBM.

Recently, highly specific and potent BH3-mimetics have been developed to specifically target BCL-2, BCL-xL and MCL-1 (12, 31, 32). We used these to probe the individual dependencies of GBM in a panel of patient-derived GSC. Importantly, we found that GBM cells are largely dependent on BCL-xL or MCL-1 for survival, whereas BCL-2 plays a lesser role. Genetic deletion of *MCL-1* corroborated its key role in both promotion and maintenance of GBM. Consistent with our findings, indirect targeting of MCL-1 through CDK7 inhibition, causing transcriptional repression, sensitises GBM cell lines to ABT-263 (47). Further, we could demonstrate that both BCL-xL and MCL-1 are highly expressed, not only in GBM tumour cores, but also in GBM stem-like cells compared to their isogenic differentiated counterparts and astrocytes. The high expression and dependence of GBM on anti-apoptotic BCL-2 function is consistent with an increased state of apoptotic priming. As Sarosiek and colleagues have demonstrated, the tissue of origin plays a major role in determining the apoptotic sensitivity of a cell (28), therefore the high dependency of stem-like cells on BCL- xL and MCL-1 might relate to their resemblance of cerebral precursor cells. Both anti-apoptotic proteins play a major role in neurogenesis, while with brain maturation neurons become refractory to apoptotic cell death (26, 28).

Unexpectedly, we observed that in comparison to their differentiated counterparts, stem-like cells were more susceptible to BH3-mimetic treatment. Investigating the basis of the differential apoptotic priming we identified NTRK2, a stem-cell specific surface receptor (42) as a key component. NTRK2 signalling, mediated by the soluble brain derived neurotrophic factor BDNF, plays a major role in cell survival promotion of growth in glial tumours (42, 48). Surprisingly, we found that BDNF stimulation led to increased apoptotic priming. This result reinforces the notion that stem-like cells are especially dependent on the tight regulation of apoptotic sensitivity. More profound understanding of the tumour-environmental context should shed light on how these interactions can be therapeutically exploited to maximise treatment efficacy of BH3-mimetics.

To facilitate translation to the clinic, we developed an *ex vivo* assay to investigate chemosensitivity of fresh patient derived GBM tissue to BCL-2 targeting BH3-mimetics. Across different freshly resected *IDHwt* GBM samples, we found that targeting both MCL-1 and BCL-xL led to an extensive induction of apoptosis and sustained reduction in tumour cell viability *ex vivo*, without compromising tissue and vessel integrity. Following inhibition of BCL-xL we found an increased amount of BIM bound to MCL-1, leading to a sensitisation of the GBM cells to MCL-1 antagonists. This mirrors studies in haematologic malignancies where the susceptibility to BH3-mimetics was dependent on BCL-2 complexed to BIM (49, 50) and increased BIM levels sensitised to BCL-2 inhibition (51).

To circumvent reported systemic toxicity, we developed a sequential treatment schedule. Our analysis of bioavailability in an orthotopic patient-derived xenograft revealed that the blood-brain-barrier is barely penetrable for ABT-263. Using the GBM cell line U87MG in a subcutaneous xenograft model, the observed effect of sequential priming could also be recapitulated *in vivo* with a profound regression of tumour size and significant survival benefit. Importantly, the brain, due to its blood-brain-barrier, provides unique opportunities for drug delivery. For instance, local drug delivery or intrathecal chemotherapy can be exerted to use the blood-brain-barrier and in turn circumvent systemic side effects (45). In summary, these data provide a rationale for further investigating alternating inhibition of BCL-xL and MCL-1 pro-survival function in GBM to maximise the therapeutic effect.

## Materials and methods

### Patient-derived GBM cell lines and cell culture reagents

Patient-derived GBM stem-like cells (E2, G1, G7, R9, R15, R24 GSC), obtained from surgical resection specimens of anonymised patients as described (52, 53), were kindly provided by Prof. Colin Watts. GSC and U87MG were cultured in serum-free Advanced Dulbecco’s modified Eagle’s medium F12 (Thermo Fisher Scientific), supplemented with 2mM glutamine, 4µg/ml heparin (Sigma), 1% B27 (Thermo Fisher Scientific), 0.5% N2 (Thermo Fisher Scientific), 20ng/ml EGF and 10ng/ml FGF (Thermo Fisher Scientific). DIFF cells were cultured in 10% foetal calf serum (FCS) containing high-glucose DMEM (Thermo Fisher Scientific) complemented with 2mM glutamine. Human astrocytes were originally provided by Prof. Steven Pollard as human foetal neural stem cells and previously differentiated to astrocytes by 7 days culture in differentiation inducing medium as described before (54). All cells were kept in 37°C incubator at 5% CO_2_ and, when grown as monolayers on Matrigel (Corning) pre-coated plates or as spheres in uncoated plates. For all experiments, cells were used up to ten passages after thawing. All cell lines used were routinely tested for mycoplasma.

For our *in vitro* studies the following drugs and chemicals were used: ABT-199 (AdooQ BioScience, #A12500-50), ABT-263 (ApexBio, #A3007), ABT-737 (ApexBio, #A8193), A1331852 and A1155463 (ApexBio, #B6164 and #B6163), S63845 (Chemgood, #C-1370), Actinomycin D (Calbiochem, #114666), q-VD-OPh (QVD, AdooQ BioScience, #A14915-25), Sytox Green (Thermo Fisher Scientific, #S7020), Brain-derived neurotrophic factor (BDNF; Peprotech, #450-02), 7,8-Dihydroxyflavone hydrate (7,8-DHF; Merck, #D5446).

### Lentiviral transduction

GSC transduction was performed using CRISPR/Cas9 genome editing with the following guide sequences:

*hBAX: 5’-AGTAGAAAAGGGCGACAACC-3’*

*hBAK: 5’-GCCATGCTGGTAGACGTGTA-3’*

*hMCL-1.1: 5’-GGGTAGTGACCCGTCCGTAC-3’*

*hMCL-1.2: 5’-GTATCACAGACGTTCTCGTA-3’*

*hBIM: 5’-* TACCCATTGCACTGAGATAG-*3’*

For stable cell line generation HEK293-FT cells (4×10^6^ in a 10 cm dish) were transfected using 4µg polyethylenimine (PEI, Polysciences) per µg plasmid DNA with the LentiCRISPRv2-puro (Addgene #52961) or LentiCRISPRv2-blasti (55) backbone, lentiviral transfer vector plasmid, packaging plasmid (Addgene #14887) and envelope plasmid pUVSVG (Addgene #8454), mixed in a 4:2:1 ratio. DNA/PEI mixtures were incubated at room temperature for 10 to 15 minutes, prior to application on HEK293-FTs. 24 and 48 hours later, virus containing supernatant was harvested and filtered (0.45µM). Virus was extracted using Lenti-X concentrator (Clontech Takara) according to the manufacturer’s instructions. The virus containing pellet was resuspended in serum-free stem-cell medium and target cells were infected in the presence of 1µg/ml polybrene (Sigma Aldrich). Two days following infection, cells were selected by growth in puromycin (E2: 1µg/ml, G7: 0.5µg/ml; Sigma Aldrich) or blasticidin (G7, R15 and R24: 10 µg/ml; InvivoGen) containing medium. As described previously, plasmids encoding iRFP IRES puro have been inserted into a pBABE vector (36).

### Cell proliferation and live-cell viability assay

Cell death and cell confluence were determined using live-cell imaging in the IncuCyte Zoom and S3 (Sartorius). For cell confluence 50×10^3^ cells were seeded in Matrigel-coated 6-well plates. Cell area per well was measured using IncuCyte imaging analysis software (Sartorius). For cell death assays, 6×10^3^ or 12×10^3^ GSC per well were seeded in Matrigel- coated 96-well plates and treated with the indicated drugs in the presence of 30nM SYTOX Green. Plates were applied to the IncuCyte imager at 37°C in a humidified 95% air/ 5% CO_2_ incubator. Every hour, two images per well were taken over a period of 24 to 48 hours. Images were presented in green phase contrast at 10x magnification. For image quantification IncuCyte imaging analysis software was used. Percentage cell death was calculated by normalising against maximal cell death control upon 24 to 48 hours treatment (1µM Actinomycin D, 10µM ABT-737 and 1µM S63845). Alternatively, 100% cell death control was verified by visual inspection of IncuCyte images, where 100% Sytox Green positive cells = total cell count.

### Clonogenic survival assay

GSC were seeded at a density of 250 cells per well in Matrigel-coated, 6-well plates with three technical repeats per experiment and left to adhere overnight. After 16 hours cells were treated as indicated for 24 hours, followed by replacement of fresh media. Cells were left to form colonies for 2 to 3 weeks prior to methanol fixation and crystal violet staining. Visible colonies consisting of minimum 50 cells were counted manually.

### Neurosphere formation assay

G7 GSC vector^CRISPR^ and MCL-1^CRISPR^ were seeded at a density of 10 cells per well in uncoated 96-well plates. Serum-free stem-cell medium was refreshed every week. Spheres were left to grow for 14 days before manual scoring of the 60 inner wells.

### Immunoblotting, immunoprecipitation and antibodies

GSC were lysed and collected in RIPA buffer (50mM Tris-HCl pH 7.5, 150 mM NaCl, 1 mM EDTA, 1% NP-40), supplemented with complete protease inhibitor (Roche) and PhosSTOP (Roche). Protein concentration was determined using Pierce BCA protein assay kit (Thermo Fisher Scientific) and protein lysates were subjected to electrophoresis through SDS–PAGE or 4-12% NuPage Bis-Tris protein gels (Thermo Fisher Scientific) followed by blotting onto nitrocellulose membranes. After blocking in 5% non-fat, dry milk or 2% BSA (Roche), membranes were probed with primary antibody (dilution 1:1000) BAK (Cell Signaling #12105), BAX (Cell Signaling #2772), BCL-2 (Cell Signaling #2762), BCL-xL (Cell Signaling #2762), MCL-1 (Cell Signaling #5453), BIM (Cell Signaling #2933), TrkB (Cell Signaling #4603), pTrkA (Tyr674/675)/pTrkB (Tyr706/707) (Cell Signaling #4621), ERK1/2 (Cell Signaling #4695), pERK1/2 (Cell Signaling #4370), AKT (Cell Signaling #9272), pAKT (Ser473; Cell Signaling #4066), Caspase 3 (Cell Signaling #9662), cleaved Caspase 3 (Cell Signaling #9664), PARP1 (Cell Signaling #9532) and SOX2 (Abcam #ab92494), NESTIN (Abcam #ab22035), GFAP (Santa Cruz #SC-6170) at 4°C overnight in blocking buffer. a- tubulin (Sigma #T5168, 1:5000), β-tubulin (Cell Signaling #2146, 1:5000), HSP60 (Cell Signaling #4870, 1:1000), or actin (Sigma #A4700, 1:5000) served as loading controls. Each blot was probed with primary antibodies and a loading control. Representative loading controls are shown in figures. Membranes were incubated with Li-Cor secondary antibodies (IRDye 680RD donkey anti-mouse, IRDye 800CW donkey anti-rabbit, IRDye 800CW donkey anti-goat) for 1 hour at room temperature.

For immunoprecipitation (IP), rabbit antibodies were coupled to magnetic beads conjugated to anti-rabbit IgG (Dynabeads Sheep anti-rabbit IgG, Invitrogen, #11203D). The buffer containing 200 mM NaCl, 75 mM Tris-HCl pH 7, 15 mM NaF, 1.5 mM Na3VO4, 7.5 mM EDTA, 7.5 mM EGTA, 0.15% (v/v) Tween-20 and protein inhibitors (Thermo Fisher) were used to prepare cell lysates. Lysates were passed several times through a 26-gauge needle followed by centrifugation at 10,000g for 5 min at 4 °C. Lysates were added to the beads and rotated for 2 hours at 4 °C. After washes in Tween-20-containing buffer, lysates were analysed by immunoblotting. Blots were imaged using Li-Cor Odyssey CLx (Li-Cor), acquired and processed using Image-Studio software (Li-Cor) and subsequently arranged using Adobe Illustrator.

### Orthotopic intracranial and subcutaneous xenografts

All mouse experiments were carried out in accordance with the Animals Act 1986 (Scientific Procedures on living animals) and the regulatory guidelines of the EU Directive 2010 under project licences PPL P4A277133 and PP6345023 and ethical review (University of Glasgow). For intracranial xenograft 7-week old female CD1-nude mice (Charles River, UK) were orthotopically injected with 1 x 10^5^ iRFP-labelled vector^CRISPR^ and MCL-1^CRISPR^ G7 GSC into the right striatum. Mice were monitored for the duration of the experiment and humanely sacrificed when they showed neurological (hemiparesis, paraplegia) or general symptoms (hunched posture, reduced mobility, and/or weight loss >20%).

For subcutaneous xenograft 1 x 10^6^ U87MG cells, previously cultured in stem-cell medium, were diluted in PBS and 50% growth factor reduced Matrigel and injected in the right flank of 8-week old female CD1-nude mice (Charles River, UK). For *in vivo* dosing, ABT-263 (ChemieTek #A263) was dissolved in 10% ethanol, 30% PEG glycol 400 and 60% Phosal 50 PG at 20 mg/kg and administered via oral gavage. S63845 (ChemieTek #S63845) was prepared in 2% vitamin E/d-a-tocopheryl polyethylene glycol 1000 succinate (Sigma) immediately prior to IV administration by tail vein injection at 25 mg/kg. Mice were treated with a 48-hours pause between drug administrations over a 14 days period. Tumour growth was monitored by caliper measurement three times per week and volume calculated using the equation ([length x width^2^]/2). Clinical end point, at which mice were euthanised, was 15mm diameter or ulceration of the tumour.

### Intravital cranial iRFP imaging and magnetic resonance imaging (MRI)

To examine intravital, intracranial tumour growth in animals bearing iRFP-positive G7 GSC, mice were monitored by PEARL imaging (Li-Cor) as previously described (36). MRI scans performed on brain tumour bearing mice using a nanoScan PET/MRI scanner (Mediso Medical Imaging Systems, Hungary). Mice were maintained under inhaled isoflurane anaesthesia (induction 5% v/v; maintenance 1.5 - 2.0% v/v) in the medical air during imaging procedure duration. Whole brain T2 Fast Spin Echo (FSE) 3D Axial Sequences (slice thickness 1.0 mm, repetition time (TR) 2000 msec, echo time (TE) 83.7 msec, Flip Angle 90 degrees) were used to acquire MRI scans. For assessments of scans, volume-of-interest (VOI) was manually drawn around the tumour region on MRI scans by visual inspection. Separate VOI were drawn for each scan to adjust for the position of the mice on the scanner and tumour size.

### Patient-derived GBM specimens and tissue culture

GBM specimens were obtained from surplus tumour tissue resected from patients treated within the OPARATIC study (NCT01390571). Patients had consented for use of surplus tissue for future research projects.

Fresh GBM tissues were obtained from surplus surgical resection tissue from patients at the Queen Elizabeth University Hospital (QEUH) in Glasgow after review by neuropathology with appropriate consent and in accordance with the NHS GG&C ethical committee review (Biorepository Application No. 432). The patient study was conducted in accordance with the Declaration of Helsinki. Neuropathological diagnosis and selected patient information are displayed in the figures. Further, details of these patients are restricted by institutional requirements. All experiments were performed conform to relevant regulatory standards of the CRUK Beatson Institute. Fresh samples were attenuated in 2% low gelling temperature agarose (Merck) and cut into 350µm thick slices using the McIlwain tissue slicer (Campden Instruments). Tissue slices were dissected under the microscope in ice cold PBS before they were transferred on top of hydrophilic Millicell cell culture inserts (Merck Millipore) into serum-free Advanced Dulbecco’s modified Eagle’s medium F12 supplemented with 0.5% N2, 1% B27, 1% glutamine and 1% penicillin-streptomycin and left to equilibrate for 24 hours at 37°C in a humidified 95% air/ 5% CO_2_ incubator, before treatment with indicated drugs for 72 hours. Following PBS washes brains were fixed in 4% paraformaldehyde (PFA) overnight.

### Organotypic adult mouse brain slice culture

Extracted brains from three 11-week old C57BL/6J mice were transferred to sterile PBS on ice, divided into both hemispheres and cut into coronal, 100µm thick slices using a vibratome (Campden Instruments 5100mz, advance speed 1mm/sec, oscillation amplitude 1.5mm, 80Hz). Up to 5 slices per hemisphere were cut around the subventricular zone (SVZ). Slices were cultured on top of cell culture inserts in neurobasal medium as described in the previous section and left to equilibrate for 1 hour at 37°C in a humidified 95% air/ 5% CO_2_ incubator before treatment with indicated drugs for 24 hours. After PBS washes slices were fixed in 4% PFA overnight.

### Immunohistochemistry (IHC) and immunofluorescence (IF)

H&E staining and IHC was performed on 4µm formalin fixed paraffin embedded (FFPE) sections. For BCL-xL (Cell Signalling #2764), cleaved Caspase 3 (Cell Signalling #9661) and MCL-1 (Abcam # ab32087) IHC staining the Leica Bond Rx Autostainer was used. All FFPE sections underwent on-board antigen retrieval for 20 minutes using ER2 retrieval buffer (Leica, UK) before staining at a previously optimised dilution (BCL-xL 1:500; cleaved Caspase 3 1:500; MCL-1 1:200) and visualised with Liquid DAB (Agilent, UK). Ki67 (Agilent #M7240) staining was performed on a Dako Autostainer Link 48 using high pH TRS retrieval buffer performed in a PT module (20 mins at 97°C). Ki67 was applied at 1:100 dilution before visualising using Liquid DAB. For SOX2, IHC epitope retrieval was achieved by heating to 98°C in pH6 citrate buffer for 60 minutes before proceeding as per the manufacturer’s instructions with SOX2 antibody used at a dilution of 1:500. Scanning and image analysis was conducted using Halo (Indica Labs). Algorithms were optimised for each stain individually and automated, quantitative analysis undertaken with Halo software (Indica Labs).

For IF staining tissue slices were permeabilised and blocked in PBS with 10% NGS, 1% BSA, 0.3% TX-100 and 0.05% Azide for 1 hour at room temperature. After washes with 10% NGS, 1% BSA, 0.1% TX-100 and 0.05% Azide containing buffer slices were incubated with primary antibodies (NESTIN 1:300, GFAP 1:400, cl. Caspase 3 1:400) in washing buffer for 72 hours at 4°C. After washes slices were incubated in secondary antibodies (1:200, Alexa Fluor 568 goat anti-mouse (#A11004), Alexa Fluor 488 goat anti-rabbit (#A11034), Life Technologies) in washing buffer for 24 hours. Following washes in PBS tissues were counterstained with DAPI (VECTASHIELD, LSBio) and mounted in gaskets (BioRad Seal Frame Incubation Chambers) on glass cover slips. Images were acquired using a Zeiss 710 laser scanning microscope with an EC Plan-Neofluar 40x/1.30 Oil DIC M27 objective and Zen 2.3 SP1 FP3 (black edition) software. 70µm Z-stacks were acquired at 2.5µm intervals and Maximum Intensity Projections (MIPs) were generated using Zen 2.1 (blue edition). Image processing was performed using Fiji (ImageJ 1.53c). Cleaved Caspase 3 positive cells were counted manually and nuclei were counted automatically using CellProfiler (Version 4.0.7).

### In silico and transcriptomic analysis

REMBRANDT microarray data was obtained from gliovis.bioinfo.cnio.es. Data was filtered for histology and tumour grading.

RNA sequencing data was obtained from a previously published GBM database (30). In order to determine the most differentially expressed genes, calculation of expression rank product was employed to assess relative gene expression in paired GSC and DIFF cell lines (56). Only results > 10 reads were incorporated.

### Statistical analyses

For comparison of two experimental groups two tailed, unpaired *t* test with Welch’s correction (Welch’s test) or Mann-Whitney test were used. For tumour related Kaplan–Meier survival curves Mantel-Cox (Log-rank) was plotted. All statistical analyses were executed with Prism software version 9 (GraphPad, La Jolla, CA, USA).

## Acknowledgements

First and foremost, our special gratitude goes to the GBM patients who agreed to make their tumour specimens available for research. Special thanks to Mary Fraser, Dr. Alexandru Stan, Dr. Zoltan Hanzely and the Neuropathology Department of the QUEH in Glasgow, who obtained patient consent and provided support with GBM tissue sampling. Further, we would like to thank the Core Services at the Cancer Research UK Beatson Institute with particular thanks to Gemma Thomson, David Strachan, the Biological Services Unit and Histology. We thank Prof. Colin Watts for generous sharing of the patient-derived GBM cell lines, Prof. Steven Pollard and Rodrigo Gutierrez Quintana for providing the human astrocytes. Many thanks to Dr. Jayanthi Anand and Dr. Dimitris Athineos for their invaluable support with the animal work. Many thanks to members of the Tait laboratory for discussion and input, and Catherine Winchester for critical reading and assistance in the preparation of this manuscript. Figures were created with BioRender.com.

## Funding

This work was supported by CRUK core funding to the Beatson Institute (A17196) to C.N., (A17196/A31287) to N.R.P., (A22903) to K.M.R. and J.O’P., and (A18277) to J.C.N., a CRUK Programme Foundation Award (C40872/A20145) to S.W.T., CRUK Clinical Research Fellowship (A23220) to A.L.K., funding by the University of Glasgow to D.K. and A.J.C. and funding by the Beatson Cancer Charity and Cancer Research UK RadNet Centre Glasgow (A28803) to K.S.. L.M.-E. was funded by the by Erasmus+ Program and Short Research stay fellowship for trainees by Universitat Autònoma de Barcelona.

## Author Contributions

A.L.K. and S.W.G.T. conceived the study, designed the work plan and wrote the manuscript. A.L.K. performed the majority of the experiments. D.K., L.M.-E., J.O’P. and F.J.B. acquired data and provided technical support. N.R.P. performed IF imaging and analysis. D.K., K.S., C.C., C.N. and G.M. provided assistance with the *in vivo* models and related imaging. K.K. and W.S. supervised and supported patient specimen collection. A.J.C. provided the RNAseq dataset. Y.E., D.K. and M.R.J. performed and supported data analysis. Expertise and critical input as well as review of the manuscript was given by K.J.C., G.I., K.B., A.J.C., K.M.R. and J.C.N.

## Competing interests

K.J.C. is a recipient of a share in royalty payments paid to the Walter and Eliza Hall Institute of Medical Research in relation to venetoclax.

---

**Supplementary Figure 1.**
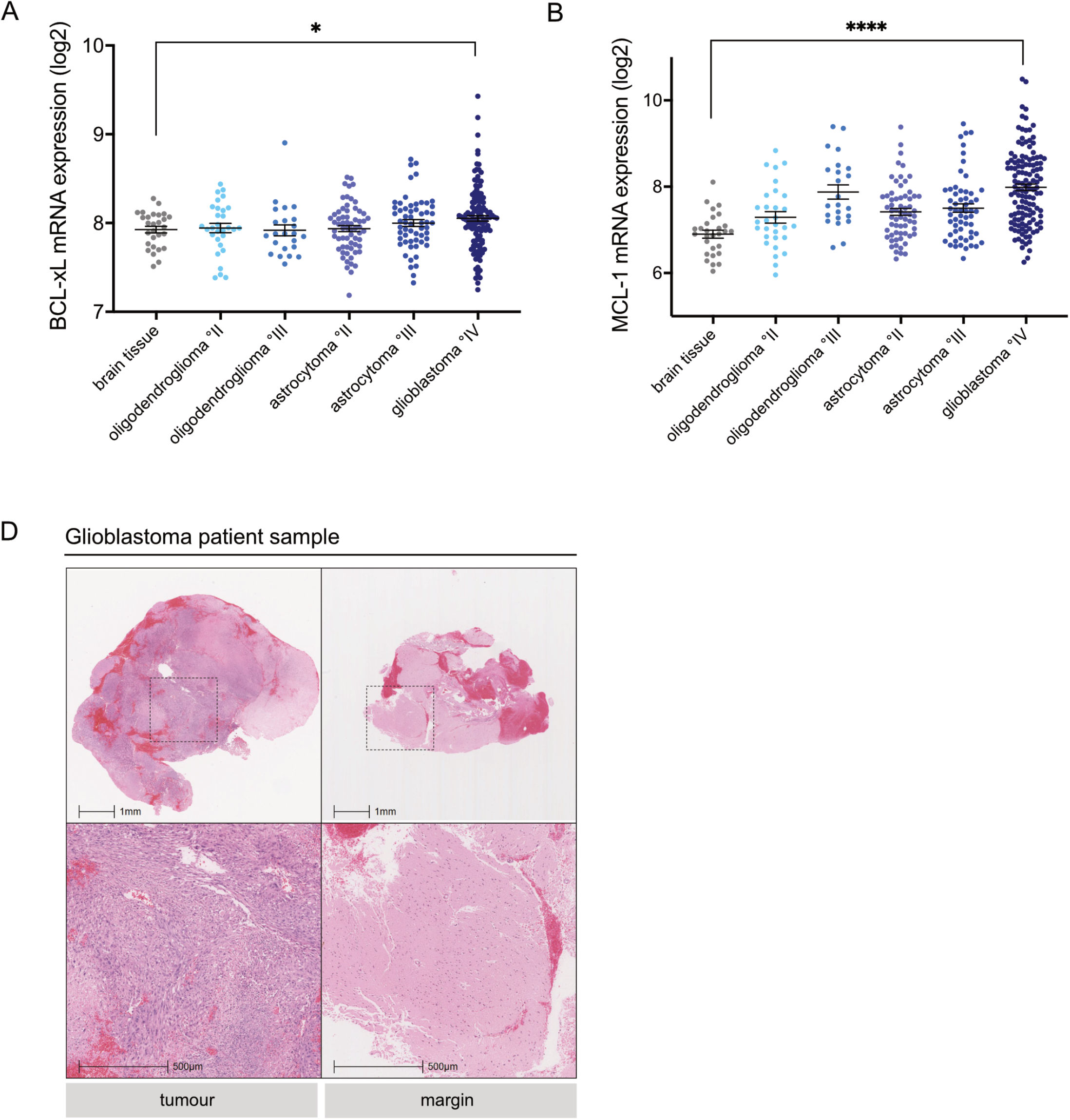
Relevant to Figure 1. (**A**,**B**) BCL-xL and MCL-1 mRNA expression from the publicly available REMBRANDT GBM microarray dataset. Data plotted for different glioma subtypes and normal brain tissue. Error bars represent mean +/-SEM (*p=0.0118, ****p<0.0001) Welch’s test. (**C**) Corresponding representative H&E images of GBM tumour and margin samples shown in lower magnification than in Figure 1D.

**Supplementary Figure 2.**
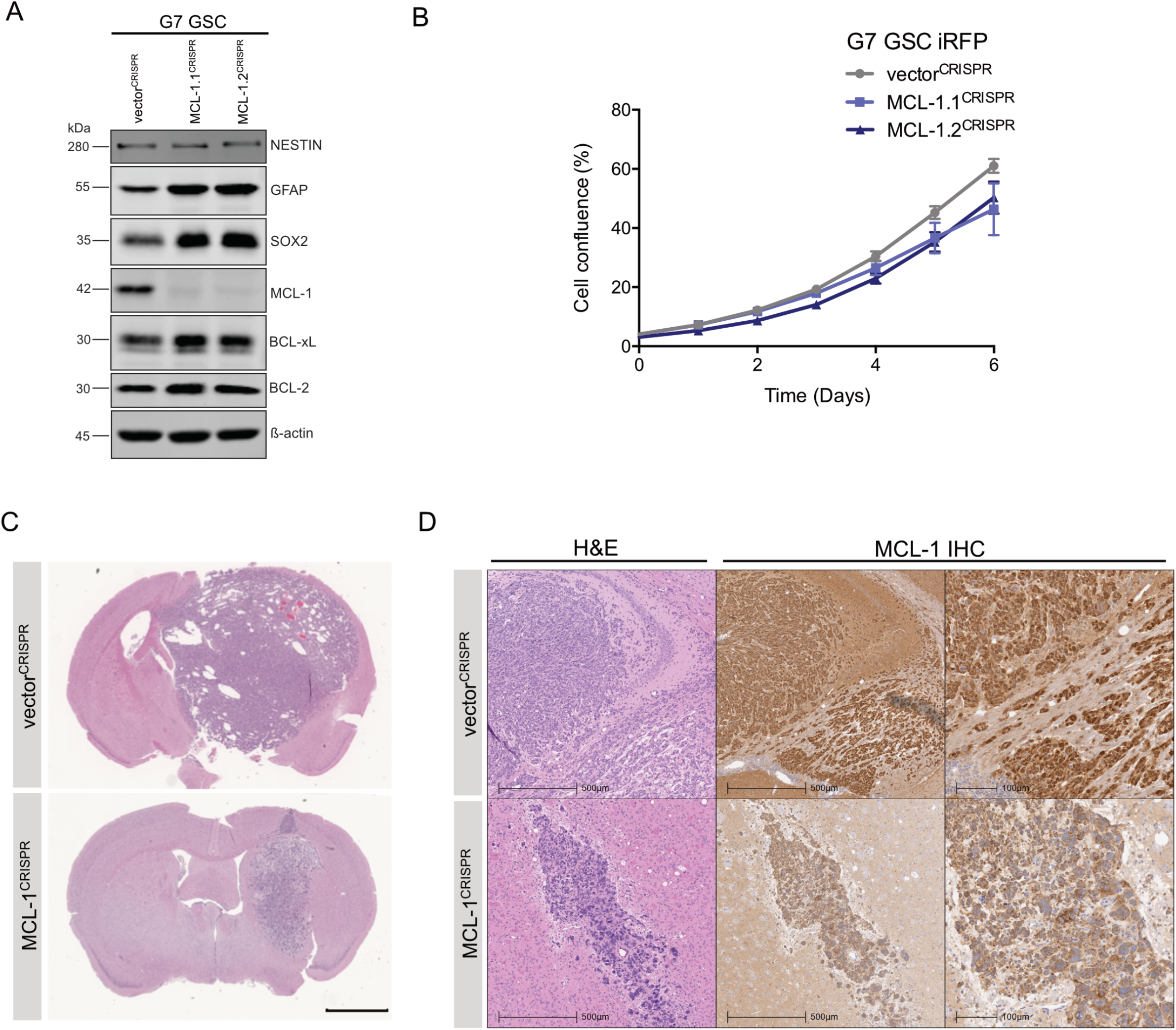
Relevant to Figure 2. **(A)** Immunoblotting of G7 GSC vector^CRISPR^, MCL1.1^CRISPR^ and MCL1.2^CRISPR^ for BCL-2 family proteins, cell-line specific neural stem cell marker (SOX2, NESTIN) and astrocyte lineage differentiation marker GFAP. Actin served as loading control. (**B**) Proliferation assay of indicated cell lines using IncuCyte Imager and – percentage cell density over 6 days. Error bars represent mean +/-SD from n=3 independent experiments. (**C**) Representative H&E images of G7 GSC iRFP vector^CRISPR^ and MCL1^CRISPR^ orthotopic xenografts at end point (corresponding to Figure 2B). Scale bar = 100µm. **(D)** Representative images of H&E and MCL-1 IHC of G7 GSC iRFP vector^CRISPR^ and MCL1^CRISPR^ orthotopic xenografts at end point.

**Supplementary Figure 3.**
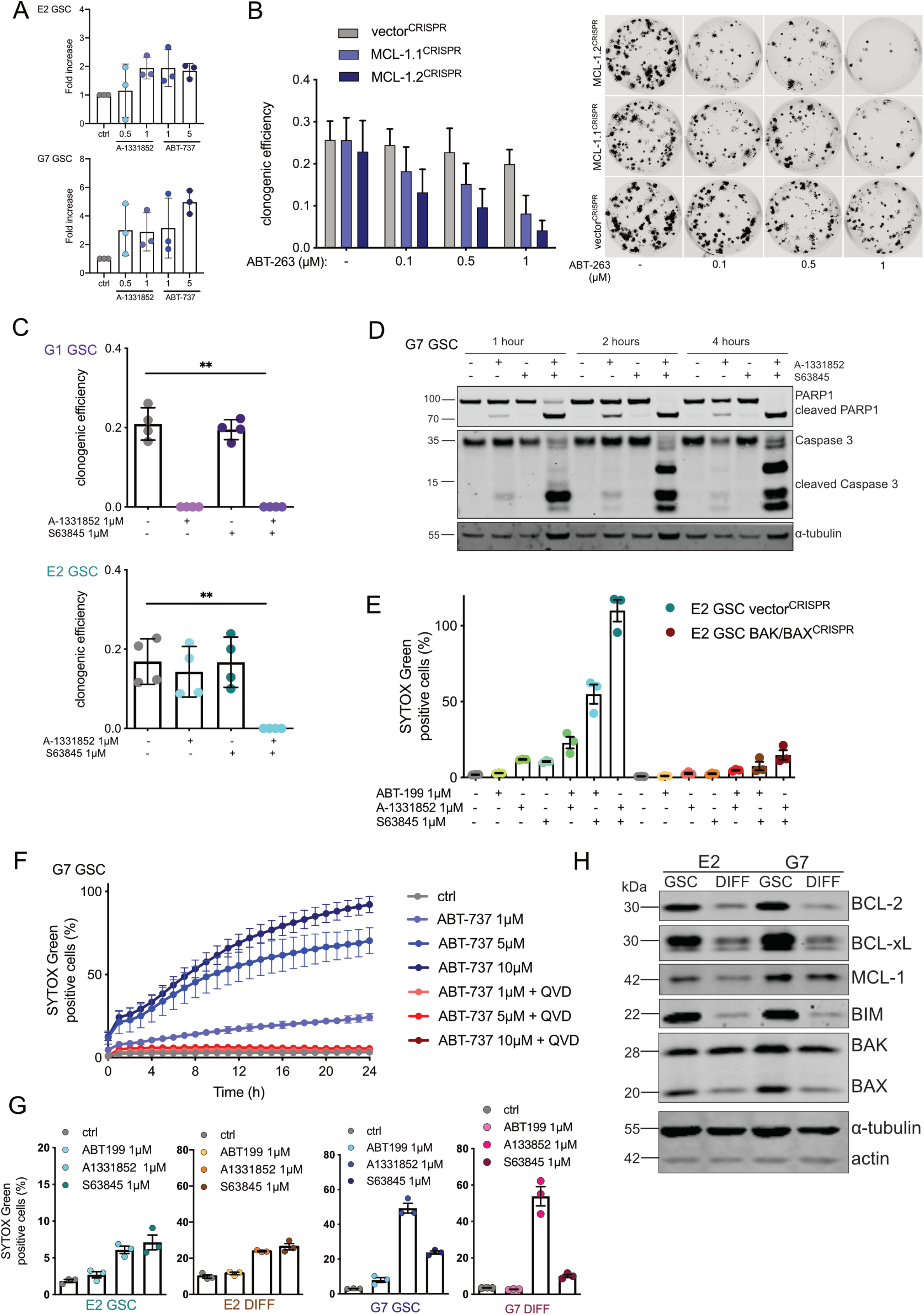
Relevant to Figure 3. **(A)** Quantification of immunoblots shown in Figure 3A. (**B**) Clonogenic survival assay of G7 GSC iRFP vector^CRISPR^ vs. MCL1.1^CRISPR^ and MCL1.2^CRISPR^ treated with indicated drugs 16 hours after plating 250 cells per well. Colonies counted manually after 14 days. Error bars represent mean +/-SD from n=4 independent experiments. Representative images scanned on LICOR imager. (**C**) Clonogenic survival assay of E2 and G1 GSC iRFP treated with indicated drugs 16 hours after plating 250 cells per well. Colonies counted manually after 14 days. Error bars represent mean +/-SEM from n=4 independent experiments (E2**p=0.0099, G1**p=0.002) Welch’s test. (**D**) G7 GSC were treated with DMSO (-), A-1331852 and S63845 for indicated times, harvested and protein expression was analysed by immunoblot. α-tubulin served as loading control. Representative image from three independent experiments. (**E**) E2 GSC vector^CRISPR^ and BAK/BAX^CRISPR^ treated with indicated drugs for 48 hours and analysed for cell viability using an IncuCyte imager and SYTOX Green exclusion. Percentage cell death was calculated by normalising against maximal cell death as described in Figure 1B. Error bars represent mean +/-SEM from n=3 independent experiments. (**F**) G7 GSC treated with indicated drugs (+/-QVD 10µM) for 24 hours and analysed for cell viability using an IncuCyte imager and SYTOX Green exclusion. Percentage cell death was calculated by normalising against maximal cell death as described in Figure 1B. Error bars represent mean +/-SEM from n=3 independent experiments. (**G**) E2 and G7 GSC with paired DIFF cells treated with indicated drugs for 24 hours and analysed for cell viability using an IncuCyte imager and SYTOX Green exclusion. Percentage cell death was calculated by normalising against maximal cell death as described in Figure 1B. Error bars represent mean +/-SEM from n=3 independent experiments. (**H**) Immunoblot of E2 and G7 GSC with paired DIFF cells for BCL-2 family proteins. α-tubulin served as loading control. Representative image from n=2 independent experiments.

**Supplementary Figure 4.**
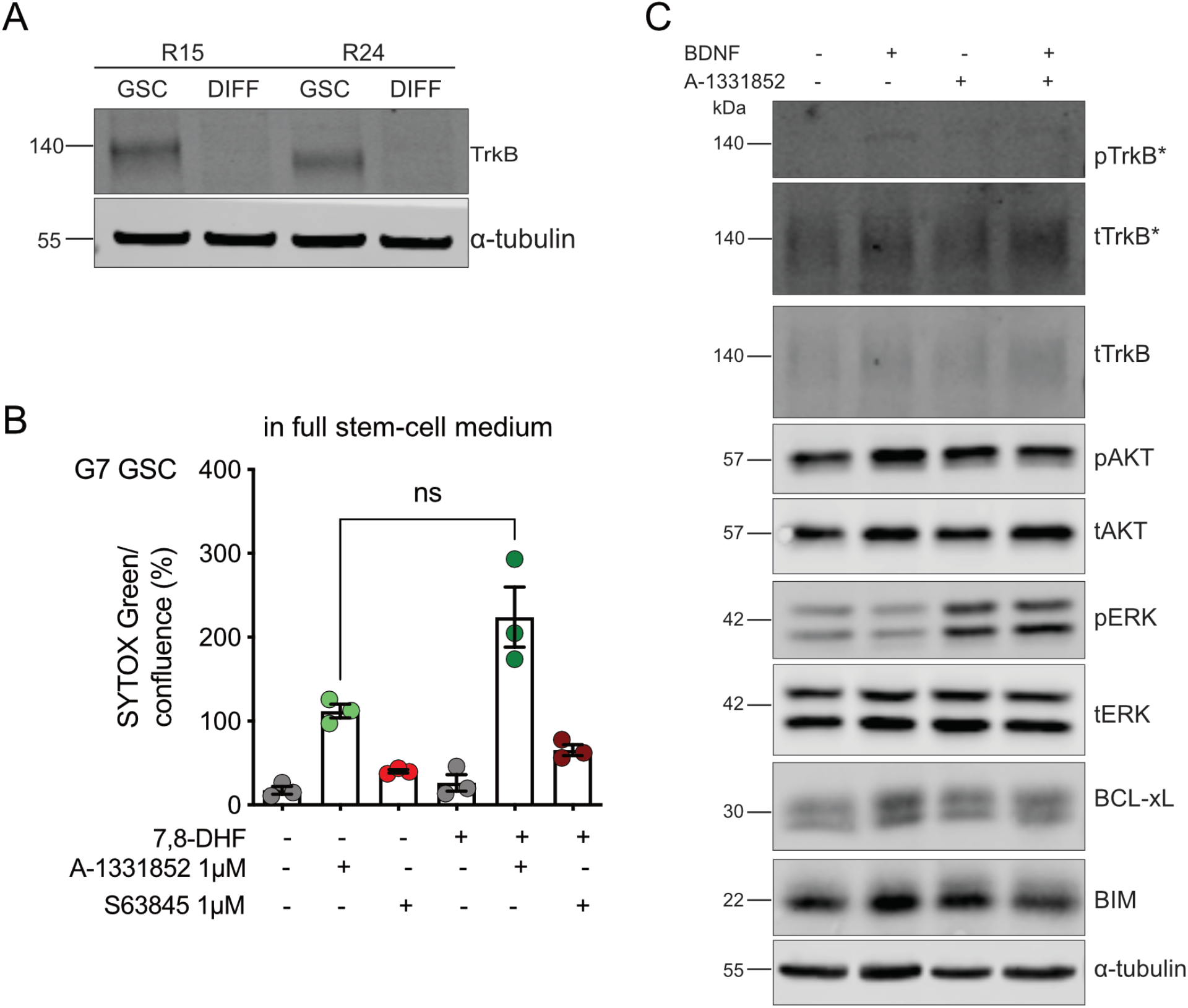
Relevant to Figure 4. **(A)** Immunoblot of TrkB in R15 and R24 GSC compared with paired DIFF cells. α-tubulin served as loading control. Representative image from n=2 independent experiments. (**B**) G7 GSC treated with indicated drugs for 24h hours in full stem-cell medium (including EGF and FGF) and analysed for cell viability using an IncuCyte imager and SYTOX Green exclusion. Error bars represent mean +/-SEM from n=3 independent experiments. (ns, p=0.0817) Welch’s test. (**C**) G7 DIFF were treated with A-1331852 2μM +/- BDNF (100ng/mL) for 1 hour, harvested and protein expression was analysed by immunoblot (*indicates high exposure). α-tubulin served as loading control. Representative image from n=3 independent experiments.

**Supplementary Figure 5.**
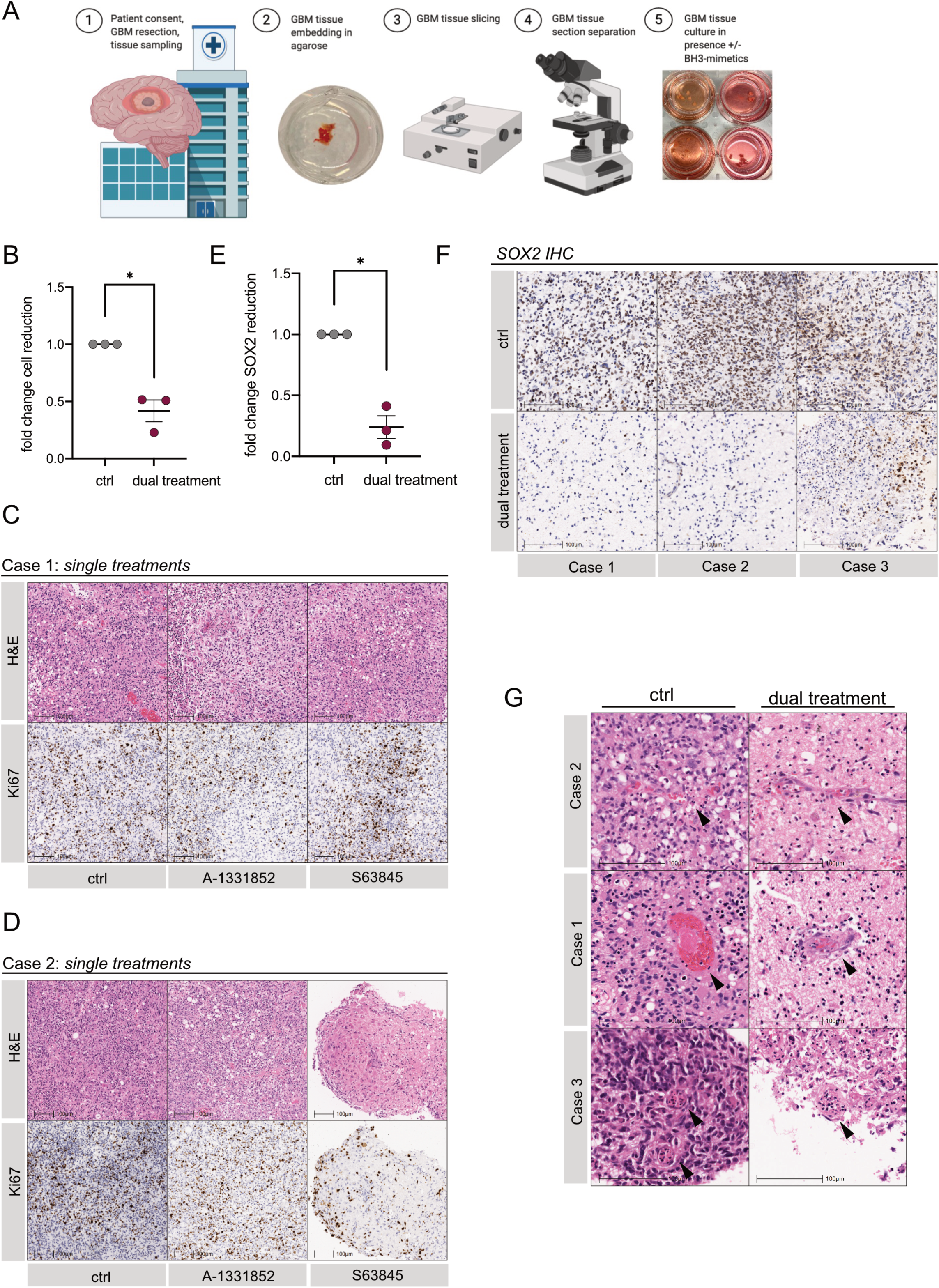
Relevant to Figure 5. **(A)** Schematic model of fresh GBM patient tissue collection and processing. (**B**) Quantification of cell reduction in GBM case 1-3 normalised to DMSO-treated control. Error bars represent mean +/-SEM (*p=0.0257) Welch’s test. (**C,D**) Representative H&E and Ki67 IHC images of GBM patients case 1 and 2 treated with single inhibitors A-1331852 2μM or S63845 2μM for 72 hours. In both cases H&E control images were used from the same area as shown in Figure 5A,B in different magnification. (**E**) Quantification of SOX2 positive cell reduction in GBM case 1-3 normalised to DMSO-treated control. Error bars represent mean +/-SEM (*p=0.0145) Welch’s test. (**F**) Representative images of SOX2 IHC in GBM case 1-3. (**G**) Representative H&E images of GBM case 1-3, arrows indicate intratumoural vessels.

**Supplementary Figure 6.** Relevant to Figure 6. (A) Immunoblot of E2 and G7 GSC vector^CRISPR^ and BIM^CRISPR^ for BCL-xL, MCL-1 and BIM. Representative images from n=2 independent experiments. α-tubulin served as loading control. (B) G7 GSC vector^CRISPR^ and BIM^CRISPR^ treated with DMSO or indicated combination for 24 hours and analysed for cell viability using an IncuCyte imager and SYTOX Green exclusion. Error bars represent mean +/-SD from n=3 independent experiments. (C) Representative images of U87MG neurospheres in stem-cell medium under treatment with indicated drugs over a total of 60 hours. Quantification of neurosphere treatment for cell viability using an IncuCyte imager and SYTOX Green exclusion given as mean signal intensity (GCU). One representative of n=3 independent experiments shown. Error bars represent mean +/-SD. (D) Clonogenic survival assay of U87MG. Treatment commenced 16 hours after plating 250 cells/well with either DMSO or ABT-263 5μM for 24 hours followed by drug washout and treatment pause for 24 hours and S63845 2μM for 24 hours. Alternating treatment was continued over the experimental period of 14 days. Colonies were counted manually. Quantification of one representative of n=2 independent experiments shown. Representative images of a replicate in one independent repeat scanned on LICOR imager. (E) Schematic of treatment schedule for *in vivo* study. (F) Percent weight change of mice (vehicle n=6, ABT-236, S63845 n=7) during the drug treatment period of the experiment in Figure 1G,H. (G) Representative H&E and Ki67 IHC images of U87MG xenografts treated with vehicle or alternating ABT-263 and S63845 therapy at end point.

## Notes

### Competing Interest Statement

The authors have declared no competing interest.

